# Frequency of mosaicism points towards mutation-prone early cleavage cell divisions in cattle

**DOI:** 10.1101/079863

**Authors:** Chad Harland, Carole Charlier, Latifa Karim, Nadine Cambisano, Manon Deckers, Myriam Mni, Erik Mullaart, Wouter Coppieters, Michel Georges

**Author notes:** Contributed equally to this work (Chad Harland, Carole Charlier).

## Abstract

It has recently become possible to directly estimate the germ-line de novo mutation (*dnm*) rate by sequencing the whole genome of father-mother-offspring trios, and this has been conducted in human^1–5^, chimpanzee^6^, mice^7^, birds^8^ and fish^9^. In these studies *dnm*’s are typically defined as variants that are heterozygous in the offspring while being absent in both parents. They are assumed to have occurred in the germ-line of one of the parents and to have been transmitted to the offspring via the sperm cell or oocyte. This definition assumes that detectable mosaïcism in the parent in which the mutation occurred is negligible. However, instances of detectable mosaïcism or premeiotic clusters are well documented in humans and other organisms, including ruminants^10–12^. We herein take advantage of cattle pedigrees to show that as much as ∼30% to ∼50% of *dnm*’s present in a gamete may occur during the early cleavage cell divisions in males and females, respectively, resulting in frequent detectable mosaïcism and a high rate of sharing of multiple *dnm*’s between siblings. This should be taken into account to accurately estimate the mutation rate in cattle and other species.

---

To study the process of *dnm*’s in the cattle germ-line, we sequenced the whole genome of 54 animals from four pedigrees. Grand-parents, parents and offspring (referred to as probands) were sequenced at average 26-fold depth (min = 21), and grand-offspring at average 21-fold depth (min = 10). The source of DNA was venous blood for females and sperm for males. The genome of one male proband (Pr 1) was sequenced both from semen (26-fold depth) and blood DNA (37-fold depth) (Figure 1A).

**Figure 1.**
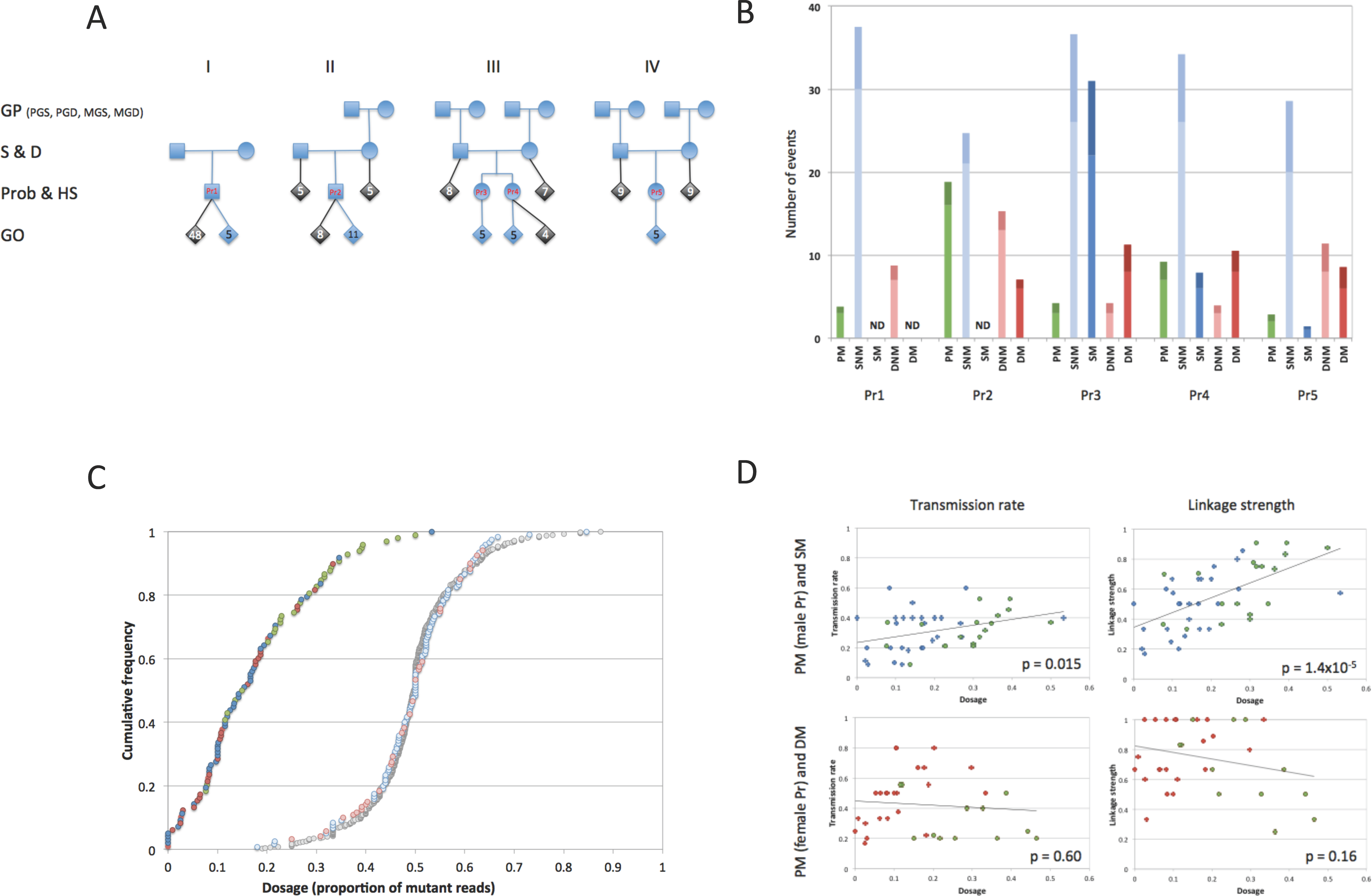
**(A) Four pedigrees (I, II, III, IV) used for the detection of *dnm*’s.** GP: grand-parents (PGS: paternal grand-sires, PGD, paternal grand-dams, MGS: maternal grand-sires, MGD: maternal grand-dams), S: sires, D: dams, Pr: probands, HS: half-sibs (of the proband), GO: grand-offspring. The five probands are labeled in red. Animals in blue were genome-sequenced at average depth of 23 and used for the detection of *dnm*’s. Animals in gray were used for confirmation by whole genome (average sequence depth of 20) or targeted sequencing (see Supplemental Methods). DNA was extracted from venous blood for females, and semen from males, except for Proband 1 for which both semen and blood DNA were analyzed. **(B) Numbers and types of *dnm*’s detected in the five probands (Pr1, Pr2, Pr3, Pr4), Pr5).** Green: PM = proband mosaic, Light blue: SNM = sire non mosaic, Dark blue: SM = sire mosaic, Light red: DNM = dam non mosaic, Dark red: DM = dam mosaic. For each bar, the lower light section corresponds to the actual number of detected *dnm*’s, the upper darker section to an extrapolation to the whole genome based on the estimated coverage. ND = not done (because the corresponding grand-parents were not sequenced). **(C) Cumulative frequency distribution of *dnm* dosage estimated as the proportion of reads carrying the mutation.** Green circles: PM mutations in the probands. Light blue circles: SNM mutations in the probands. Light red circles: DNM mutations in the probands. Dark blue circles: SM mutations in the sires. Dark red circles: DM mutations in the dams. Grey circles: corresponding PM, SNM, DNM, SM and DM mutations in the grand-offspring. The three SM and one DM variant with dosage of 0, were shared between the proband and at least one half-sib yet not detectable in the semen or blood of the corresponding parent. **(D)** Relationship between the *dnm* dosage (fraction of mutant reads) and the rate of transmission to offspring (left) and strength of linkage (right) for mutations that are detectably mosaic in the sperm of a male parent (upper), or in the blood of a female parent (lower). Green circles: PM mutations, blue circles: SM mutations, red circles: DM mutations. The corresponding correlations were significant in males (p = 0.015 and 1.4×10^−5^) but not in females (p = 0.60 and 0.16).

Using the standard definition, we identified 190 candidate *dnm*’s as variants that were (i) detected in a proband, (ii) absent in both parents (and grand-parents when available), (iii) transmitted to at least one grand-offspring, and (iv) not previously reported in unrelated individuals from the 1,000 Bulls project^13^ (Suppl. Figure 1&2 and Suppl. Table 1). For confirmation, we developed amplicons spanning 113 candidate *dnm*’s and sequenced them at average depth of ∼2,187 in the 54 animals plus 55 relatives (Figure 1A). This confirmed the genuine nature of 110/113 variants, demonstrating the excellent specificity of our bioinformatics pipeline. The three remaining ones were also detected in one of the parents (although not in the grand-parents) in the confirmation, and momentarily ignored.

**Table 1:** Relative likelihood of the observations under different models of gametogenesis. The first four columns correspond to the parameters that were tested in the model: (i) the fold increase of the mutation rate (1×, 10×, 20×), (ii) during the x first cell divisions (4, 7, 11, 15, 18), (iii) the number of PGCs (4, 10, 40), and (iv) the ontogenetic relatedness of the PGCs (F(alse) or T(rue)). Log(LR) corresponds to the logarithm (base 10) of the likelihood of the data relative to the best model (first line). Parameters are in bold when the corresponding model is the best given that parameter value. We only show results for models that are the best given at least one parameter value. Likelihoods of all models are given in Suppl. Table 3.

| Fold increase mutation rate | During first x cell divisions | Number of PGCs | Related PGCs or not | Log(LR) |
| --- | --- | --- | --- | --- |
| **20x** | **4** | **4** | **T** | 0.00 |
| **10x** | 4 | 4 | T | -0.35 |
| 20x | 4 | 4 | **F** | -0.40 |
| 10x | **7** | 4 | F | -1.57 |
| 20x | 4 | **10** | T | -1.74 |
| 20x | 4 | **40** | T | -3.45 |
| 10x | **11** | 4 | T | -5.09 |
| 20x | **15** | 4 | T | -7.03 |
| 10x | **18** | 4 | T | -9.46 |
| **1x** | 7 | 4 | T | -16.17 |

We first examined what proportion of *dnm*’s detected in a proband might actually have occurred during its development rather than being inherited via the sperm or oocyte. An unambiguous distinction between the two types of *dnm*’s is their degree of linkage with either the paternal or maternal haplotype upon transmission to the next generation (i.e. the grand-offspring in Figure 1A). *Dnm*’s that have occurred in the germ-line of the sire will show *perfect* linkage with the proband’s paternal haplotype in the grand-offspring (i.e. always transmitted with the paternal haplotype, never transmitted with the maternal haplotype), while *dnm*’s that have occurred in the germ-line of the dam will show *perfect* linkage with the proband’s maternal haplotype in the grand-offspring. On the contrary, *dnm*’s that have occurred during the development of the proband will be in *complete* (but imperfect) linkage with either the paternal or maternal haplotype (i.e. sometimes transmitted with the maternal haplotype, never transmitted with the paternal haplotype, or sometimes transmitted with the paternal haplotype, never transmitted with the maternal haplotype)(Suppl. Figure 1). Across the four pedigrees, 124 variants were in perfect linkage with the paternal haplotype, 32 in perfect linkage with the maternal haplotype, 10 in complete (but imperfect) linkage with the paternal haplotype and 21 in complete (but imperfect) linkage with the maternal haplotype (Figure 1B). If the 10+21 *dnm*’s indeed occurred during the development of the proband rather then being inherited from the sire or dam, the *dnm* dosage (defined as the proportion of reads spanning the *dnm* site that carry the mutant allele) is expected to be < 50% in the proband but equal to 50% in the grand-offspring inheriting the *dnm*. The mean dosage was 0.26 in the proband, and 0.52 in the grand-offspring, and this difference was highly significant (p < 10^−6^). The corresponding means were 0.48 and 0.49 (p = 0.40) for the 124+32 mutations showing perfect linkage with either the paternal or maternal haplotype (Figure 1C).

We conclude that in cattle ∼17% of *dnm*’s detected in an animal using standard procedures are not inherited from the sire or dam but correspond to premeiotic clusters generated during the development of the individual. This is a lower bound, as *dnm* will only be detected and recognized as having occurred in the proband if (i) the *dnm* dosage is sufficiently high for the proband to be called heterozygote, (ii) the *dnm* is transmitted to at least one grand-offspring, and (iii) complete (but imperfect) linkage is demonstrated in the grand-offspring (Suppl. Figure 3). We will refer to this type of *dnm*’s as Proband-Mosaic (PM), while the others will be referred to as Sire-Non-Mosaic (SNM) (meaning that the sire is not mosaic for a mutation transmitted via his sperm), or as Dam-Non-Mosaic (DNM) (meaning that the dam is not mosaic for a mutation transmitted via her oocyte). The proportion of PM (but not SNM and DNM) mutations differed significantly between probands (p = 0.004); neither differed significantly between sexes (p > 0.30). Of particular interest, the three PM mutations of proband 1 were detected in both sperm and blood DNA (Suppl. Table 1), indicating that they occurred early in development (see hereafter).

If detectable mosaicism for *dnm*’s is common in the individual in whom they occurred, requiring their absence in the DNA of the parents (as typically done) will cause genuine *dnm*’s to be eliminated. We took advantage of the grandparents available in three pedigrees to recover such events as variants that were (i) absent in the grand-parents, (ii) detected in either sire or dam with a dosage significantly < 50% (Suppl. Table 1), (iii) transmitted to the proband with a dosage of ∼50%, (iv) transmitted to at least one grand-offspring with a dosage of ∼50%, and (v) not previously reported in unrelated individuals^13^. We will refer to these types of mutations as Sire-Mosaic (SM) and Dam-Mosaic (DM), respectively (meaning that the sire/dam is detectably mosaic for a *dnm* transmitted via the sperm/oocyte)(Suppl. Figure 1). We detected 61 such candidate events, including the 3/113 variants mentioned above (Suppl. Table 1 and Suppl. Figure 2). We developed amplicons for 34, and sequenced (average 1,498-fold depth) all 54 individuals plus 55 relatives (including ≥ 5 half-sibs of the probands) (Figure 1A). We took advantage of whole genome sequence information that became available for 27 half-sibs, to trace the inheritance of the remaining 24 candidate variants. The ensuing data indicated that 11/61 candidates were genuine *dnm*’s but occurred in the germ-line of one of the grand-parents rather than one of the parents (dosage ∼50% in the sire or dam, and perfect linkage in the half-sibs). The SM/DM status was unambiguously demonstrated for 40 (dosage < 50% in the sire or dam in the confirmation, transmission to half-sibs, and complete (but imperfect) linkage) and strongly supported for the remaining 10 (dosage < 50% in the sire/dam in the confirmation or complete (but imperfect) linkage yet without transmission) (Suppl. Figure 2 and Figure 1B). Further supporting the genuine nature of the SM/DM mutations, the dosage was 0.12 on average in the corresponding parent, while being 0.51 in descendants (p < 10^−6^) (Figure 1C).

**Figure 2:**
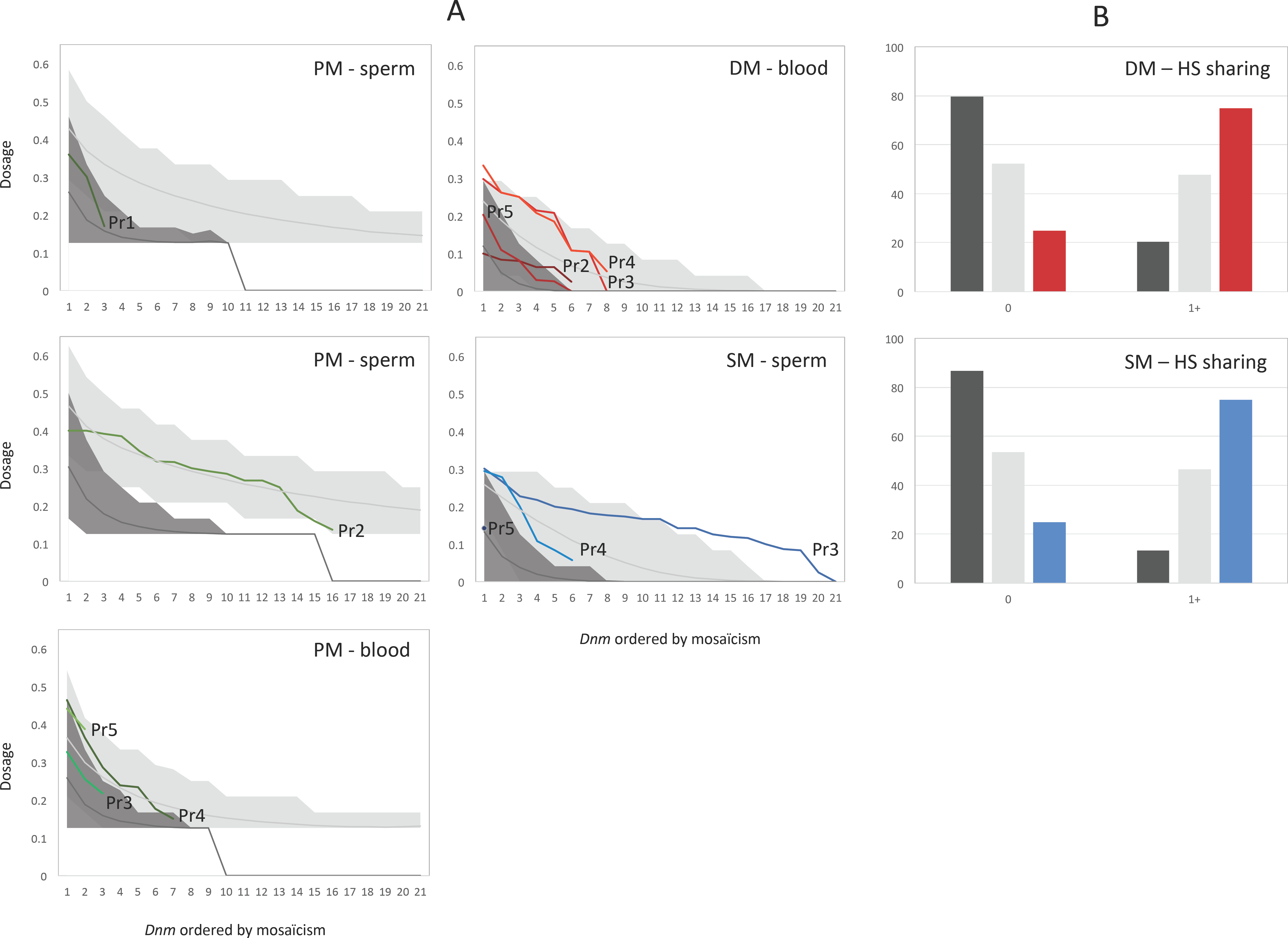
**(A)** *Dnm*’s with detectable mosaicism in sperm DNA of male probands (PM – sperm) or sires (SM – sperm), or in blood DNA of female probands (PM – blood) or dams (DM – blood), ranked by observed rate of mosaicism. Colored lines: real data. Pr1-5: proband 1-5. Dark gray shaded area: 95% confidence interval obtained from simulations assuming uniform mutation rate per cell division and 40 unrelated PGCs. Light gray shaded area: 95% confidence interval obtained from simulations assuming 20-fold higher mutation rate during the first 4 cell divisions, and 4 related PGCs (Table 1). **(B)** Distribution of the proportion of half-sibs (HS) of the probands that share 0, or at least 1 (1+) of the *dnm*’s detected in the corresponding proband. Red bars: real observations for *dnm’*s transmitted by the dam (DM+DNM). Blue bars: real observations for *dnm’*s transmitted by the sire (SM+SNM). Dark grey bars: expectation under the null hypothesis of uniform prenatal mutation rate per cell division and 40 unrelated PGCs. Light grey bars: expectation under the best alternative model assuming a 20× increased mutation rate during the first 4 cell division and 4 related PGCs (Table 1).

For *dnm*’s that were detectably mosaic in sperm (SM and PM in male probands), allelic dosage was significantly correlated with rate of transmission (p = 0.025) and strength of linkage (p = 0.0002). These correlations were not significant for *dnm*’s that were detectably mosaic in blood (DM and PM in female probands). This suggests that the degree of mosaicism in the soma is a poor indicator of the degree of mosaicism in the germ line (Figure 1D). Accordingly, the rate of transmission of the three PM mutations of proband 1 to its 53 offspring was better predicted by their dosage in sperm than in blood (Suppl. Figure 4).

Considering SNM/SM and DNM/DM mutations jointly, we conclude that on average a sire is detectably mosaic (in sperm) for 29% of *dnm*’s present in a sperm cell, while a dam is detectably mosaic (in blood) for 51% of *dnm*’s present in an oocyte. These are lower bounds as we only considered *dnm*’s for which the dosage was significantly < 0.5 in the parent (condition (ii) above). These figures are possibly consistent with recent reports in the mouse (∼25%)^7^, but considerably larger than current estimates in human (∼5%)^14^. They are certainly larger than expected assuming that the mutation rate per cell division is uniform throughout development, and it suggests that the mutation rate is higher for early cell divisions (Suppl. Figure 5). Moreover, when analyzing the transmission patterns of SM and DM mutations to the half-sibs of the proband (in whom the *dnm*’s were detected), we were struck by the fact that (i) >60% of half-sibs share at least one *dnm* with the proband, while <50% are expected (p = 0.05), and (ii) half-sibs sharing multiple *dnm*’s with the proband appeared surprisingly common (Suppl. Table 2). These findings also indicate that a substantial proportion of *dnm*’s must occur early in development and be present in the precursor cells common to the soma and germ line (Suppl. Figure 5).

In mammals, after fertilization, cleavage, and segregation of (i) the inner cell mass from the trophoblast, (ii) the epiblast from the hypoblast, (iii) the embryonic epiblast from the amniotic ectoderm, a small number of epiblast-derived cells located in the wall of the yolk sac in the vicinity of the allantois are induced to become primordial germ cells (PGCs). These migrate to the primitive gonad where they expand and produce >1 million gametogonia. Oogonia initiate meiosis prior to birth in females. Spermatogonia will resume mitotic divisions at puberty allowing (i) the maintenance of a pool of stem cell like spermatogonia, and (ii) sustained spermiogenesis involving ∼3 additional mitotic divisions followed by meiosis (Suppl. Figure 6). We simulated the process of de novo mutagenesis in the male and female germ cell lineages assuming (i) uniform pre- and post-natal mutation rates per cell division, and (ii) 40 PGCs sampled at random from the embryonic epiblast-derived cells^15^. Pre- and post-natal mutation rates were adjusted to match the observed number of mutations per gamete (34 in sperm, 14 in oocytes). Under these conditions, we virtually never observed the level of mosaicism, nor the sharing between sibs characterizing the real data (Figure 2). We (i) increased the relative mutation rate during the early cell divisions (keeping the mutation rate per gamete constant)(10 and 20-fold increase during the first 4, 7, 11, 15 and 18 cell divisions; Suppl. Figure 6), (ii) reduced the number of induced PGCs (4, 10, or 40), and (iii) varied the relatedness between PGCs (i.e. sampled randomly amongst all embryonic epiblast-derived cells or from a sub-sector)(Suppl. Figure 6). Increasing the mutation rate during the very first cell divisions matched the real data much better (Figure 2). To quantitatively evaluate model fitting we used (i) the proportion of PM, SM and DM mutations with corresponding rate of mosaicism in sperm and soma, and (ii) the proportion of sibs sharing 0, 1, 2,…*dnm*’s with a proband, to compute the likelihood of the data under different scenarios (see M&M). A 20-fold increased mutation rate during the first four cell divisions, combined with 4 related PGCs fitted the data best (Table 1 and Suppl. Table 2). The data were ≥10^16^ times less likely under models assuming a uniform mutation rate throughout development, and ≥10^5^ times less likely assuming an increased mutation rate passed the 7^th^ cell division (after segregation of inner cell mass and throphoblast)(Table 1 and Suppl. Table 2).

When accounting for genome coverage, the estimated number of *dnm*’s per gamete (SNM+SM, DNM+DM) averaged 46.6 for sperm cells and 18.1 for oocytes (male/female ratio of 2.6), corresponding to an average mutation rate of ∼1.2×10^−8^ per base pair per gamete. Including an estimate (from the simulations) of the number of missed SM (∼3.3) / DM (∼1.1) and misclassified PM mutations (∼2.9 to ∼8 depending on the proband), yields an average mutation rate of ∼1.17×10^−8^ per base pair per gamete and a male/female ratio of 2.4. The standard approach of ascertaining *dnm*’s (i.e. erroneously considering PM mutations, ignoring SM and DM mutations) would have yielded a mutation rate of 0.9×10^−8^ per bp per gamete, with a 2.5-fold higher mutation rate in bulls than in cows.

Two hundred twenty of the 237 identified *dnm*’s were nucleotide substitutions, 17 small insertion-deletions. The non-mosaic classes of mutations (SNM and DNM) were ∼30-fold enriched in CpG>TpG transitions as expected. This signature was also present but less pronounced for mosaic mutations (PM, SM and DM). Mosaic mutations were ∼2.6-fold enriched in C>A and/or G>T transversions, largely due to GpCpA>GpApA and TpCpT>TpApT substitutions (Figure 3). This was unlikely to be an artifact for reasons spelled out in Suppl. Figure 7. It is noteworthy that this is exactly the same mutational signature as the one recently reported for human embryonic somatic mutations^16^. There was no obvious difference between the profile of *dnm*’s in the male and female germ line (data not shown). In general, *dnm*’s appeared uniformly scattered across the genome (Suppl. Figure 8).

**Figure 3:**
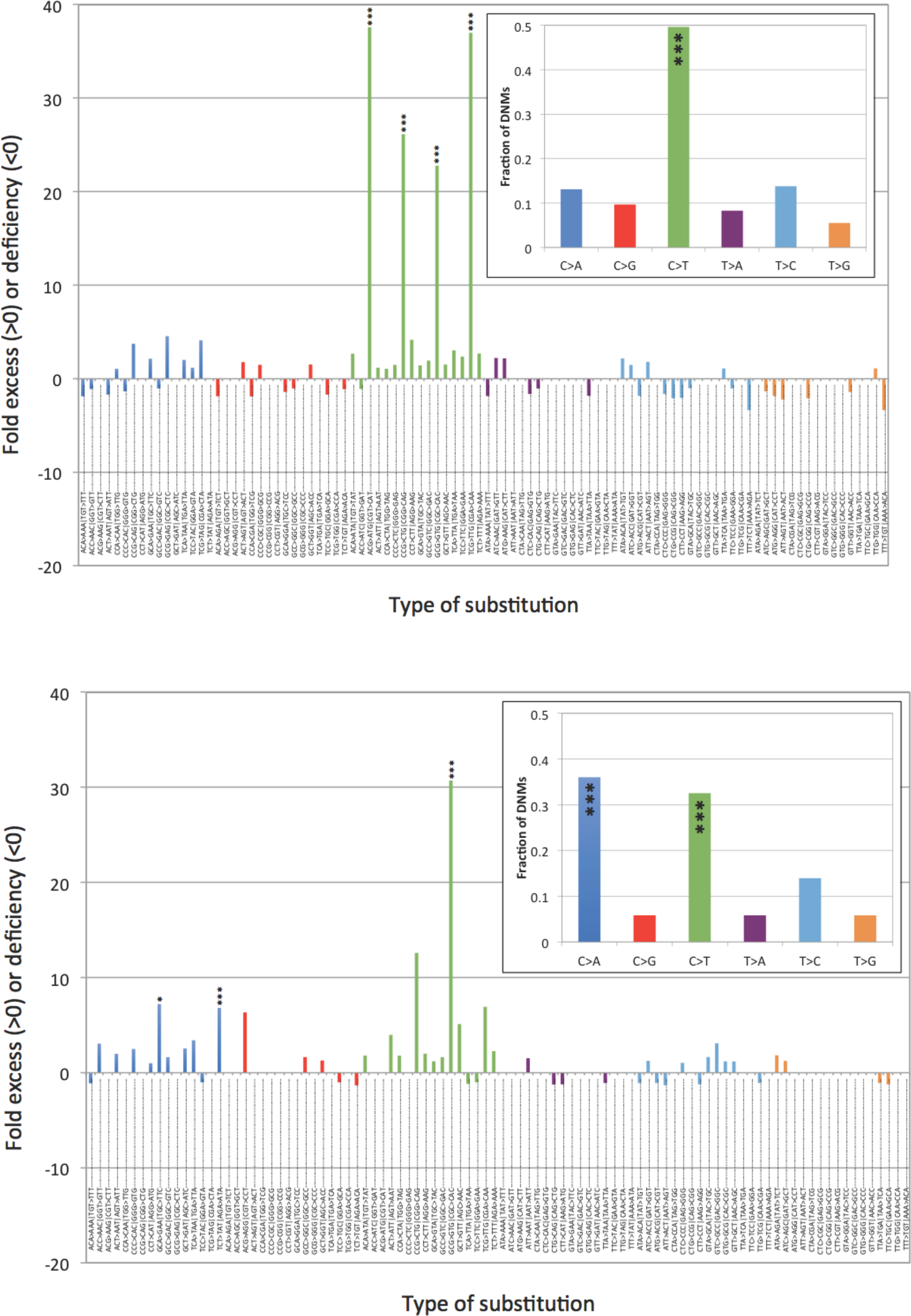
**(A)** SNM and DNM (i.e. *dnm*’s assumed to have occurred in the later stages of gametogenesis): fold excess or deficiency over expected for specific nucleotide substitutions accounting for trinucleotide context. Inset: Proportion of *dnm*’s corresponding to the six possible types of nucleotide substitutions. **(D)** Idem for PM, SM and DM (i.e. *dnm*’s assumed to have occurred in the early stages of gametogenesis). ***: p< 0.001, *: p< 0.05 (accounting for multiple testing by Sidak correction).

The enrichment of C>A/G>T transversions in the mosaic mutations caused the overall Ti/Tv ratio to be 1.33, well below expectations. This was likely due to sampling variation (meaning that Ti/Tv ratios might differ between families and that we by chance sampled families at the low end), as the Ti/Tv ratio was 1.99 for 2,530 candidate *dnm*’s detected with the same bioinformatics pipeline in a follow-up study of 113 probands (excluding the ones analyzed in this work), i.e. closer to expectations and the 2.2 Ti/Tv ratio of SNPs segregating in the Holstein-Friesian dairy cattle population (MAF ≤ 0.01; rare allele considered to be the derived allele). However, the spectrum of the 2,530 *dnm*’s remained significantly different from the SNP spectrum, with an excess of C>A/G>T transversions in the mosaic class of mutations, an excess of C>T/G>A transitions in both mosaic and non-mosaic mutations, and a paucity of T>C/A>G transitions in both mosaic and non-mosaic mutations (Suppl. Figure 7). This could point towards recent alterations of the mutational profile in domestic cattle. It is worth noting in this regard that most analyzed animals were bred using artificial insemination and/or in vitro embryo production. It seems unlikely that artificial insemination with frozen semen could explain the observed familial clustering of specific *dnm*’s. However, it is conceivable that in vitro maturation, fertilization and culture of oocytes and embryos affect the *dnm* rate, possibly by perturbing DNA replication. It is important to determine whether this is the case, especially as the same methods are increasingly used in human reproduction.

*Dnm*’s occurring during the development of an individual, should a priori affect the maternal and paternal chromosome with equal probability. When considering the PM, SM and DM jointly, 48 mosaic mutations occurred on the maternal chromosome versus 31 on the paternal chromosome (p = 0.11). This trend suggests that the maternal and paternal chromosomes might be epigenetically distinct during early development and that this may affect their mutability.

Our work points towards the fact that direct estimates of mutation rates from sequencing families may have to be revisited, taken PM, SM and DM status into account, to obtain more accurate estimates of the mutation rate per gamete and per generation. This may affect both the overall mutation rate as well as its male/female ratio. However, our analyses suggest that the effect is likely to be modest and would, for instance, be insufficient to explain the present 2-fold discrepancy between direct and indirect estimates in human studies^17,18^. We confirmed by simulation that the rate of mosaicism does not significantly affect the rate of nucleotide substitution per generation or average fixation time^19^ (Suppl. Figure 9).

Our work calls for a careful reevaluation of the importance of mosaicism for *dnm*’s in humans. If more common than presently appreciated, the recurrence risk of *dnm*-dependent disorders in sibs may be higher than generally assumed ^11,17^. Moreover, a non-negligible proportion of true *dnm*’s may have been ignored (because they were detected at low dosage in the parents) in *dnm*-dependent searches for genes underlying inherited disorders hence reducing the potential power of such studies.

## Acknowledgements

This work was funded by the DAMONA Advanced ERC project to Michel Georges. Carole Charlier is Senior Research Associate from the Fonds de la Recherche Scientifique - FNRS (F.R.S.-FNRS). Chad Harland has been funded in part by Livestock Improvement Corporation (New Zealand). We are grateful to Erik Mullaart and CRV (Arnhem the Netherlands) for providing us with the sperm and blood samples. We used the supercomputing facilities of the Consortium des Equipements de Calcul Intensif en Fédération Wallonie Bruxelles (CECI) funded by the F.R.S-FNRS.

## Authors contributions

MG, CH, CC: designed the experiments. EM: provided samples. LK, NC, MD, WC: performed the sequencing. CH, MG, CC: analyzed data. MG, CH, CC: wrote the paper.

## Data availability

All sequence data will be made freely available in public databases.

## Methods

### Whole genome sequencing

DNA was extracted from sperm (for males) or whole blood (females and one male) for the four families and their relatives using standard procedures. Familial relationships were confirmed by genotyping all DNAs with the 10K Illumina SNP chip. We constructed 550bp insert size whole genome Illumina Nextera PCR free libraries following the protocols recommended by the manufacturer. All samples where then sequenced on Illumina HighSeq 2000 instruments, using the 2×100bp paired end protocol by the GIGA Genomics platform (University of Liege). Data was mapped using BWA mem (version 0.7.9a-r786)^20^ to the BosTau6 reference genome. Alignments were processed according to the GATK^21^ best practices version 2 with PCR duplicates marked, Indel realignment and Base Quality Score Recalibration using known sites. GATK HaplotypeCaller (version 3.4) was used according to the N+1 workflow to generate variants from the alignments. Common variants were then compared to a 10K Illumina SNP chip for each individual to confirm the identity of the library.

### Detection of de novo mutations

We developed a suite of scripts to identify *dnm*’s from a vcf file produced by GATK and containing sequence information about members of four-generation pedigrees such as the ones described in Fig. 1. The first (“4_phaser_4_gen.pl”) generates the linkage phase for the parents (sire and dam), the proband, and the grand-offspring. Phasing is done based on high quality variant positions and genotypes (f.i. QUAL score ≥ 50,000; PL scores ≥ 100; sequence depth ≤ 2.5× the average sequence depth). The outcome is knowledge of the grand-parental origin of the paternal and maternal chromosomes of the proband including the identification of cross-over events, as well as the grand-parental origin of the chromosomes transmitted by the proband to the grand-offspring including the identification of cross-over events. The second module (“5_de_novo_detector_4_gen.pl”) identifies the candidate *dnm*’s per se. It first identifies diallelic variant positions for which all grand-parents, sire, dam and proband have a genotype and sequence coverage between set limits (f.i. 10 and 60). The proportion of variants sites satisfying these depth limits was used to estimate the proportion of the genome (of the total of the 2,670,422,299 base pairs in the bovine Bostau6 build) that was explored. Candidate *dnm*’s were then identified as sites for which (i) the QUAL score was ≥ 100, (ii) the proband had genotype 0/1 with corresponding PL-scores ≥ 40, (iii) none of the grand-parents had reads with the alternate allele (AD = x,0), (iv) either the sire or the dam had no reads with the alternate allele, and (v) reads with the alternate allele were found in the grand-offspring. The same script also determines the genotype frequencies (0/0, 0/1 and 1/1) at the corresponding position in sequenced individuals outside of the pedigree that are not descendants of the sire or the dam. The third module (“6_germline_assigner_4_gen.pl”) combines the output of the first and second module to determine in which individual the *dnm* is most likely to have occurred (one of the four grand-parents, sire or dam, or proband) and on which grandparental chromosome it occurred. All candidate *dnm*’s were manually curated using the Integrated Genome Viewer (IGV)^22^. *Dnm*’s with mutant reads in either sire or dam (even if called 0/0 by GATK) were relabeled as SM or DM, provided that the *dnm* segregated in complete (but imperfect) linkage with the paternal or maternal haplotype (respectively), in the half-sibs of the proband.

To test the corresponding pipeline, we identified 11,255 variants (of which 10,093 SNPs) for which Pr2 was heterozygous and which were not present in unrelated individuals including from the 1,000 Bulls project. The corresponding “genotype fields” of the parents and grand-offspring were modified in the vcf file such that the genotype (GT) was set at 0/0 and the unfiltered allele depth (AD) of the derived allele set at 0. The Ti/Tv ratio for the corresponding SNPs was 1.9922. The proportion of the genome explored was estimated at 76% as described above. The pipeline detected 9,325 variants (83% sensitivity) of which 8,409 SNPS (81% sensitivity). The Ti/Tv ratio amongst detected *dnm*’s was 1.9989, indicating that the pipeline did not introduce a Ti/Tv bias.

### Confirmation of candidate dnm’s

PCR primers were then designed for each candidate passing the quality check in IGV using BatchPrimer^23^ targeting a product size of 200-1000bp with at least one primer being present in unique (non-repeat) sequence (as identified by repeatmasker). The resulting amplicons were sequenced on an Illumina MiSeq instrument using the 2×250bp paired end protocol. The sequenced amplicons were aligned to the BosTau6 reference genome using BWA mem and candidate de novo mutations were checked in IGV and variants were called using freebayes (v1.0.2-15-g357f175)^24^.

### Modeling gametogenesis

<u>(i) Data types:</u> To compare the adequacy of the different gametogenesis models we computed the likelihood of three types of data. The first is the degree of mosaicism in the parent across *dnm*’s detected in a given gamete. Thus we may have detected *n* SM and *m* SNM *dnm*’s in a given sperm cell. The *n* SM *dnm*’s have dosages in the paternal sperm DNA of x_1_, x_2_, x_3_,…x_n_ > 0 while the *m* SNM *dnm*’s have a dosage of 0. We have three such lists for Pr3, Pr4 and Pr5. Likewise, we may have detected *n* OM and *m* ONM *dnm*’s in a given oocyte. The *n* OM *dnm*’s have dosages in the maternal blood DNA of x_1_, x_2_, x3,…x_n_ > 0 while the *m* ONM *dnm*’s have a dosage of 0. We have four such lists for Pr2, Pr3, Pr4 and Pr5.

The second data set consists in lists of PM *dnm*’s and their dosage (x_1_, x_2_, x_3_,…x_n_ > 0) in sperm (Pr1 and Pr2) or blood DNA (Pr3, Pr4, Pr5).

The third data type consists in the number of *dnm*’s detected in a gamete transmitted to a proband, and the numbers of those shared by the studied half-sibs of the proband. Thus, we may have detected *n* SM and *m* SNM *dnm*’s in a sperm cell (or oocyte) transmitted to a given proband, of which half-sib 1 will share x_1_, half-sib 2 x_2_,…,where x_i_ ≤ *n*.

<u>(ii) Computing probabilities under various models of gametogenesis:</u> For data type 1, we simulated the process of de novo mutation in the female and male cell lineages described in Suppl. Figure 6. For the null hypothesis, the mutation rate per cell division before birth was set at an average of 0.77 (Poisson distributed), such that the number of *dnm*’s per oocyte averaged 14 (as observed). The mutation rate per cell division after birth was set at an average of 0.3 (Poisson distributed), such that the number of *dnm*’s per sperm cell averaged 34 (as observed). For alternative hypotheses, the mutation rate for the early cleavage cell divisions (4, 7, 11, 15 and 18 first cell divisions, corresponding to the different development stages in Suppl. Figure 6) was increased 10- or 20-fold when compared to the remaining prenatal cell divisions, for which the mutation rate was concomitantly reduced such that the overall number of *dnm*’s per oocyte remained unaffected (average of 14). We further tested 4, 10 and 40 induced PGCs, and unrelated or related induced PGCs as described in Suppl. Fig. 6. For all 90 possible scenarios, we determined by simulation what proportion of *dnm*’s found in a sperm cell (respectively oocyte) were characterized by a dosage in paternal sperm (respectively maternal soma) of 0-0.05, 0.05-0.10,… These proportions were then used as probabilities in computing the likelihood of the data (i.e. a series *dnm*’s with corresponding rate of mosaicism in the parental tissue) under the corresponding model. Given our experimental design, SM and DM are only recognized as such, if (i) their dosage in the sire (SM) or dam (DM) is significantly < 0.5, and (ii) they show complete (but imperfect) linkage in the available half-sibs. <u>We considered a fixed number of eight half-sibs in the simulations.</u> These conditions were included in the simulations. Thus a mutation was only considered if it satisfied these two criteria. The dosage that was considered was not the true dosage for that mutation, but the “realized” dosage assuming a sequence depth of 24.

To compute the likelihood of the second type of data, we simulated gametogenesis in exactly the same way as for data type 1. We then randomly sampled *n* gametes, where *n* corresponds to the number of GO (hence 5 for Pr1, Pr3-5, and 11 for Pr2). For all *dnm*’s in these *n* gametes, we then determined the dosage in the germ-line (Pr1, Pr2) or soma (Pr3-5). For all 90 possible scenarios, we determined by simulation what proportion of PM *dnm*’s detected in sperm of blood DNA were characterized by a dosage of 0-0.05, 0.05-0.10,… These proportions were then used as probabilities in computing the likelihood of the data (i.e. a series PM *dnm*’s with corresponding dosage in sperm or blood) under the corresponding model. With the real data, PM mutations are only recognized as such (i) if they are transmitted to at least one of the *n* offspring, (ii) if the proband is called heterozygous for the corresponding *dnm* by GATK, and (iii) if we demonstrate complete (but imperfect) linkage in the GO. Condition (i) is achieved in the simulation by sampling *n* gametes at random. Condition (ii) and (iii) were modeled in the simulations. <u>We considered five and eleven GO to match the real data.</u> Thus a mutation was only considered if it satisfied these two criteria. The dosage that was considered was not the true dosage for that mutation, but the “realized” dosage assuming a sequence depth of 24.

For data type 3, we modified the simulations in order to exactly generate a predetermined number *n* of *dnm*’s in a given “reference” gamete. Thus if in the real data an oocyte was characterized by 11 *dnm*’s (f.i. Pr3 and Pr4), we would in the simulations distribute 11 *dnm*’s across the (7+4+4+3+18) cell divisions leading to a simulated reference oocyte and track their segregation (according to their point of occurrence) across the entire germ line lineage. Under the null hypothesis of uniform prenatal mutation rate, all 36 cell divisions would have equal chance to be hit by anyone of the 11 mutations. Under the alternative hypotheses, early cleavage cell divisions would have a 10- or 20-fold higher chance than the remaining ones. Under the hypothesis of related PGCs the segregation pattern of early mutations in the germ line lineage would be concomitantly affected (see Suppl. Fig. 6). We would then samples gametes at random from the same germ line tree and count the number of mutations shared with the “reference” gamete. This would generate a frequency distribution of gametes sharing 0, 1, 2,…, *n dnm*’s with the reference gamete. The corresponding frequencies were then used as probabilities in computing the likelihood of the data (i.e. a series half-sibs sharing 0, 1,…,*n dnm*’s with the reference gamete transmitted to the proband) under the corresponding model.

Likelihoods of the data under the 90 tested models were then simply computed as the product of the probabilities of all *dnm*’s (data type 1 and 2) and half-sibs (data type 3) extracted from the simulations performed under the corresponding model.

<u>(iii) Estimating the number of missed SM and DM and misclassified PM mutations:</u> The simulations for dataset 1 allowed us to estimate the number of SM and DM mutations missed either because the “realized” dosage was to high in the parent, or because we could not demonstrate complete (but imperfect) linkage in the half-sibs. Likewise the simulations for dataset 2 allowed us to estimate the number of PM mutations that, although detected (realized dosage sufficient to be called heterozygous by GATK), were misclassified as SNM or DNM mutations because showing perfect linkage in the available GO. Under the best biological model (20x increased mutation rate during the first 4 cell divisions, 4 related PGCs, see Table 1), these numbers were: (i) average loss of 3.3 SM mutations, (ii) average loss of 1.1 lost DM mutations, (iii) average gain of 1.45 SNM and 1.45 DNM mutations for a male proband with 11 GO, (iv) average gain of 4 SNM and 4 DNM mutations for a male proband with 5 GO, and (v) average gain of 1.8 SNM and 1.8 DNM mutations for a female proband with 5 GO.

